# Investigating the role of melatonin in bipolar disorder using transcriptomics

**DOI:** 10.1101/2025.09.03.673928

**Authors:** Heather K. Macpherson, Trang T. T. Truong, Michael Berk, Ken Walder, Susannah J. Tye

**Author notes:** H. K. Macpherson and T. T. T. Truong are co-first authors. **Address for correspondence:** Susannah J. Tye, PhD, Functional Neuromodulation and Novel Therapeutics Laboratory, Queensland Brain Institute, The University of Queensland – Brisbane, QLD, Australia. **Declaration of interest:** The authors declare that they have no known competing financial interests or personal relationships that could have appeared to influence the work reported in this paper. **CRediT author statement: Heather Macpherson:** Conceptualisation, Formal Analysis, Visualisation, Writing – Original draft preparation, Project administration. **Trang T. T. Truong:** Conceptualisation, Methodology, Investigation, Formal Analysis, Writing – Review and Editing. **Michael Berk:** Writing – Reviewing and Editing, Resources. **Ken Walder:** Writing – Reviewing and Editing, Resources. **Susannah Tye:** Conceptualisation, Writing – Reviewing and Editing, Supervision, Project administration.

## Abstract

Melatonin may be a potential therapeutic target for bipolar disorder (BD) treatment; however, its role in BD pathophysiology remains poorly understood. This study aimed to investigate the therapeutic and mechanistic role of melatonin in BD using transcriptomics. RNA sequencing (RNAseq) data from 216 post-mortem dorsolateral prefrontal cortex samples (156 controls, 60 BD) were used to generate gene regulatory networks (GRNs). These were compared to lists of melatonin-related genes using gene set enrichment analysis (GSEA) to assess differential expression between people with BD and controls. Furthermore, RNAseq data from NT2-N cells treated with lithium, lamotrigine, valproate, or quetiapine were compared to lists of melatonin-related genes using GSEA. Finally, BD-associated gene regulatory patterns were compared to GRNs induced by melatoninergic agents to evaluate the repurposing potential of these pharmacotherapies for BD. Genes involved in inhibiting melatonin signalling were nominally upregulated in the BD post-mortem gene expression dataset. Transcription factors (TFs) activating melatonin signalling tended to be downregulated in BD females, while TFs inhibiting melatonin signalling were significantly downregulated in BD males. Regarding current treatments, quetiapine caused the greatest number of significant alterations in melatonin-related gene expression, followed by valproate, lithium, and then lamotrigine. Valproate was found to significantly upregulate genes involved in melatonin degradation. Finally, the melatonin receptor agonist GR-135531 was identified as a possible repurposing candidate for BD. Overall, this study provides new evidence that dysregulation of melatonin-related genes may play a role in the pathophysiology of BD, and suggests a number of melatoninergic agents as potential therapeutic candidates for BD.

## 1 Introduction

Melatonin may be a therapeutic target for the treatment of bipolar disorder (BD), which is increasingly recognised as a disorder of circadian dysregulation, with disrupted sleep-wake cycles, hormonal rhythm disturbance, and affective instability across diurnal periods.^1,2^ Exogenous melatonin has shown potential efficacy in treating manic and depressive episodes in BD patients.^3–7^ People with BD in all mood states frequently demonstrate reductions in plasma melatonin, suggesting hypomelatoninaemia (low melatonin secretion and function) is a trait marker of BD, and thus may have a causal role.^8–11^ Unfortunately, neither melatonin’s mechanism of action nor the pathophysiological underpinnings of hypomelatoninaemia in BD are well understood.

Given that therapeutics that target underlying pathophysiology are more likely to improve treatment outcomes, achieve symptom resolution and prevent relapse, understanding the causal mechanisms underpinning BD may assist in finding novel, more effective treatments.^12^ Investigating the effects of currently used BD therapeutics on potential underlying causes such as melatonin dysregulation may also be beneficial for elucidating their mechanisms of action, and ultimately assist with precision psychiatry.

This study aimed to clarify the therapeutic and mechanistic role of melatonin in BD using transcriptomics. Specifically, we curated a comprehensive list of genes related to melatonin synthesis, signalling, and degradation, and assessed the differential regulation patterns of these genes between people with BD and healthy controls (HCs) using gene regulatory networks (GRNs) and gene set enrichment analysis (GSEA). We also assessed whether current BD medications altered the expression and regulation of melatonin-related genes in a human neuronal cell model. Additionally, we used GRNs to identify the effectiveness of repurposing melatonin receptor agonists to treat BD.

## 2 Methods

A summary of the methodology used in this study is demonstrated in Figure 1.

**Figure 1:**
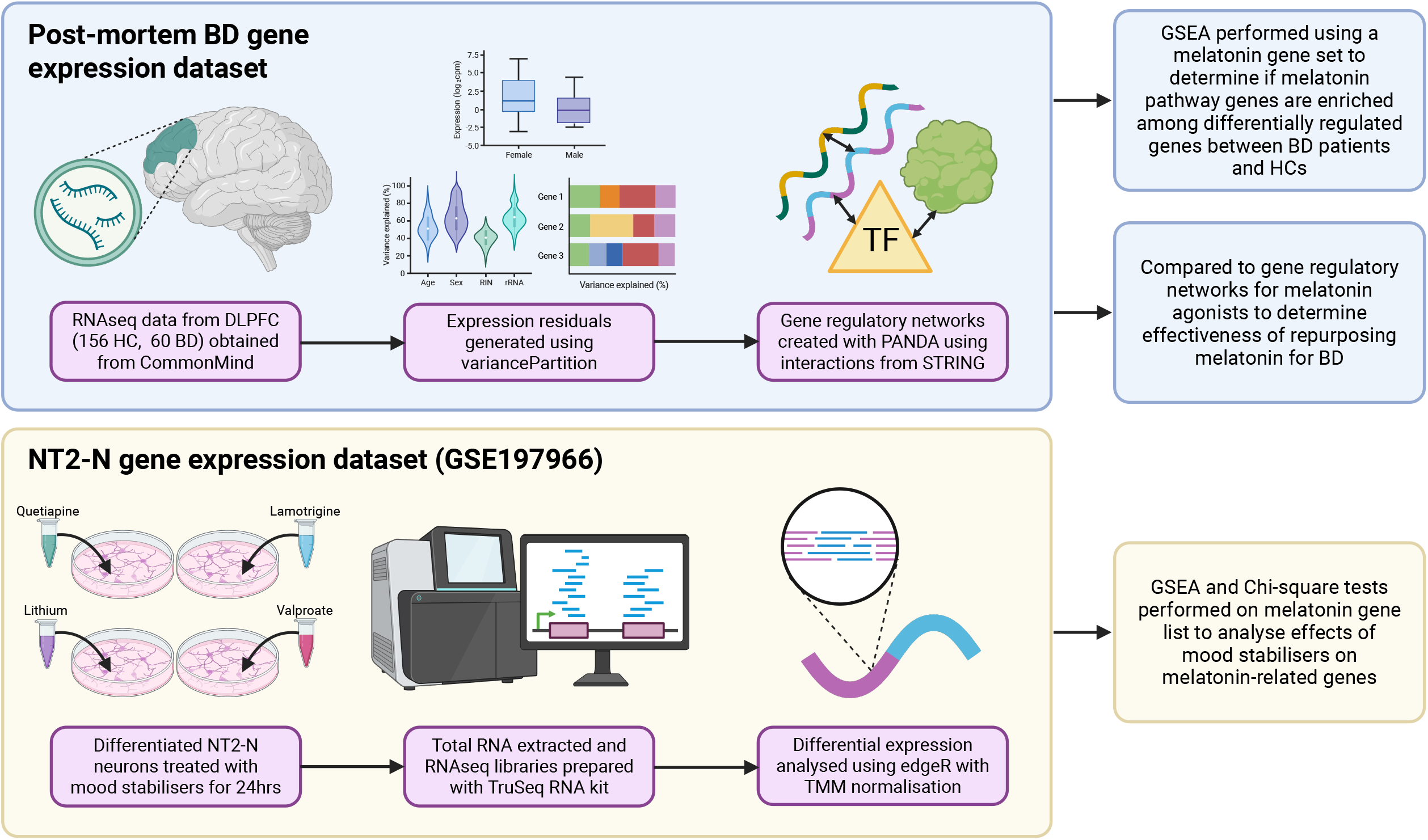
Study design. The post-mortem bipolar disorder (BD) gene expression dataset was created using RNA sequencing (RNAseq) data from post-mortem dorsolateral prefrontal cortex (DLPFC) samples obtained from BD patients and healthy controls (HC). This dataset was used to create gene regulatory networks, which were then compared to a list of genes related to melatonin synthesis, signalling, and degradation using gene set enrichment analysis (GSEA), and to gene regulatory networks for melatonin agonists. The NT2-N gene expression dataset was created using RNAseq libraries prepared from differentiated NT2-N neurons treated with various mood stabilisers for 24 hours. This dataset was then compared to a list of melatonin-related genes using GSEA. [Created by authors with biorender.com]

### 2.1 Gene sets used in this study

#### 2.1.1 Post-mortem bipolar disorder gene expression dataset

This dataset consists of RNA sequencing (RNAseq) data sampled from post-mortem tissue. The processing of this dataset and gene regulatory network (GRN) generation were conducted as previously described.^13^ Concisely, RNAseq data from 216 post-mortem dorsolateral prefrontal cortex (DLPFC) samples (156 HCs, 60 BD) from the HBCC Brain Bank were obtained via the CommonMind Consortium.^14^ Expression residuals were generated using the variancePartition R package,^15^ regressing out technical and biological covariates, and used for co-expression analysis. GRNs were constructed using PANDA,^16^ integrating gene co-expression, TF binding motifs,^17,18^ and TF-protein interactions from STRING.^19^ Networks were built separately for BD and control groups using high-confidence interactions, and filtered to retain overlapping genes and TFs.

Gender-specific analyses were conducted in a similar fashion. Samples were stratified by sex (39 BD males, 117 HC males; 21 BD females, 39 HC females), and expression residuals were generated separately for each gender cohort using variancePartition, again controlling for technical and biological covariates. Co-expression analyses were performed within each group, and PANDA was used to construct GRNs using the same integrative approach. Within each gender, networks were built separately for BD and HC groups, filtered to retain overlapping genes and TFs, and compared to assess differences between BD and controls.

#### 2.1.2 NT2-N gene set

This dataset consists of RNAseq data collected from human NT2 teratocarcinoma cells (ATCC, United States; RRID: CVCL_0034). The differentiation, treatment, extraction of RNA, and preparation of RNAseq libraries was conducted as previously described.^20^ Briefly, these cells were differentiated into post-mitotic neurons (NT2-N) using retinoic acid over 28 days, and then treated for 24 hours with lithium (2.5 mM), lamotrigine (50 µM), quetiapine (50 µM), valproate (0.5 mM), or appropriate vehicle controls. Doses were selected based on prior work demonstrating no cytotoxicity and consistent transcriptional effects.^21,22^ Total RNA was extracted post-treatment using the RNeasy^®^ Mini kit (QIAGEN, Germany; Cat: #74106). RNAseq libraries were prepared using the TruSeq RNA Sample Preparation kit (Illumina, United States; Cat: #RS-930-200) and sequenced on a HiSeq 2500 System (Illumina, United States; 50 bp single end reads). Quality control, alignment (STAR v2.5, GRCh38), and expression quantification were performed using a standardised pipeline (https://github.com/m-richardson/RNASeq_pipe). Raw data are available at the Gene Expression Omnibus database (accession number: GSE197966).

### 2.2 Melatonin gene list

A list of genes involved in melatonin synthesis, signalling, and degradation was manually curated from pathway databases. These include the Kyoto Encyclopedia of Genes and Genomes (KEGG) pathways hsa04713 (Circadian entrainment), hsa04010 (MAPK signalling pathway), hsa04710 (Circadian rhythm), map00380 (Tryptophan metabolism), hsa4150 (mTOR signalling), hsa04151 (P13K-Akt signalling) and hsa0464 (NF-kappa B signalling)^23,24^; and the Qiagen GeneGlobe pathways ‘Melatonin Signaling,’ ‘Superpathway of Melatonin Degradation,’ ‘Melatonin Degradation III,’ and ‘Circadian Rhythm Signaling’.^25^ The effects of melatonin on individual genes were confirmed using available literature.

### 2.3 Differential regulation of melatonin genes in people with bipolar disorder

A differential network approach was applied using the aforementioned post-mortem BD gene expression dataset and melatonin gene list. This approach assessed the regulation patterns of melatonin-related genes in BD through network-based differential targeting scores. Differential targeting was quantified by calculating the difference in edge weights between BD and control networks. Within each network, two types of targeting scores were calculated: a gene-based targeting score, defined as the sum of inbound edge weights, reflecting how strongly each gene is regulated by others; and a TF-based targeting score, defined as the sum of outbound edge weights, representing the regulatory influence of each TF. These scores were used as ranking metrics for gene set enrichment analysis (GSEA). GSEA was performed using ClusterProfiler on the curated melatonin-related pathways.^26^ Gene-based scores were used to assess whether melatonin-related genes were preferentially up- or downregulated in BD, while TF-based scores were used to evaluate whether melatonin-related TFs acted as key regulators in BD. Statistical significance was assessed via permutation testing with false discovery rate (FDR) correction.

### 2.4 Differential expression of melatonin genes by drugs used for bipolar disorder

The previously described NT2-N gene expression data were used to assess whether current BD medications up- or downregulate genes involved in melatonin signalling, synthesis, and/or degradation using differential expression analysis, GSEA, and Chi-squared tests. Differential expression was analysed using edgeR,^27^ with TMM normalisation and FDR correction via the Benjamini-Hochberg method. GSEA was performed using the clusterProfiler R package,^26^ incorporating the curated melatonin-related pathways. Genes were ranked by their log2 fold change values obtained from differential expression analysis.

Differential expression results for only melatonin-related genes were extracted and categorised by significance (q < 0.05 threshold) and whether melatonin signalling was opposed (i.e., “activator” genes were downregulated or “inhibitor” genes were upregulated) or enhanced (i.e., “activator” genes were upregulated or “inhibitor” genes were downregulated) by each drug. Contingency tables were created according to these categorised results, and analysed with Chi-squared tests (Supplementary Table 1).

### 2.5 Evaluating effectiveness of repurposing melatonin for bipolar disorder

GRNs for melatonin agonists were generated using the GRAND database by integrating transcriptomic perturbation data from the Connectivity Map (CMAP) with multi-omic regulatory priors.^28,29^ Melatonin receptor agonists (melatonin, ramelteon, agomelatine, and GR-135531) were identified from the small molecule compounds profiled in CMAP, which includes gene expression data across various cell lines, doses, and time points (173 013 treatment conditions in total). For each compound, a GRN was constructed by integrating TF binding motifs, protein–protein interaction data, and drug-induced gene expression profiles using the PANDA algorithm, producing a weighted bipartite GRN.^16^ To quantify regulatory influence, a targeting score was calculated for each TF as the sum of its outbound edge weights to all genes. TFs with differential targeting were identified as those with targeting scores more than two standard deviations from the mean across all compounds. Due to computational constraints, only the most representative treatment condition for each compound was retained in the final TF-by-drug targeting matrix—specifically, the condition yielding the highest number of TFs with differential targeting.

Gender-specific analyses were conducted using the same framework. Treatment conditions were stratified by donor gender of the cell lines to choose the most representative treatment condition for each compound, i.e., the condition yielding the highest number of TFs with differential targeting. Treatments from male donor samples were compared only to the male BD regulatory query, and those from female donor samples were compared only to the female BD regulatory query.

To identify melatonin agonists capable of reversing disease-associated regulatory patterns, cosine similarity between the BD regulatory query and drug-induced gene targeting profiles were applied, which compares differentially targeted gene lists between the input and each drug.^30^ A score of −1 indicates complete reversal of the input pattern (suggesting therapeutic potential), while +1 indicates reinforcement (potentially harmful).

## 3 Results

### 3.1 Melatonin gene list

Melatonin influences a wide range of pathways involved in circadian entrainment (e.g., CLOCK/BMAL1), calcium homeostasis (e.g., PLCβ/PKC/CaMKK), glucose homeostasis (e.g., PI3K/Akt/GSK3β/mTOR), inflammation (e.g., MAPK/ERK or MAPK/JNK), and oxidative stress (Akt/FOXO3/SIRT1) (Figure 2).

**Figure 2:**
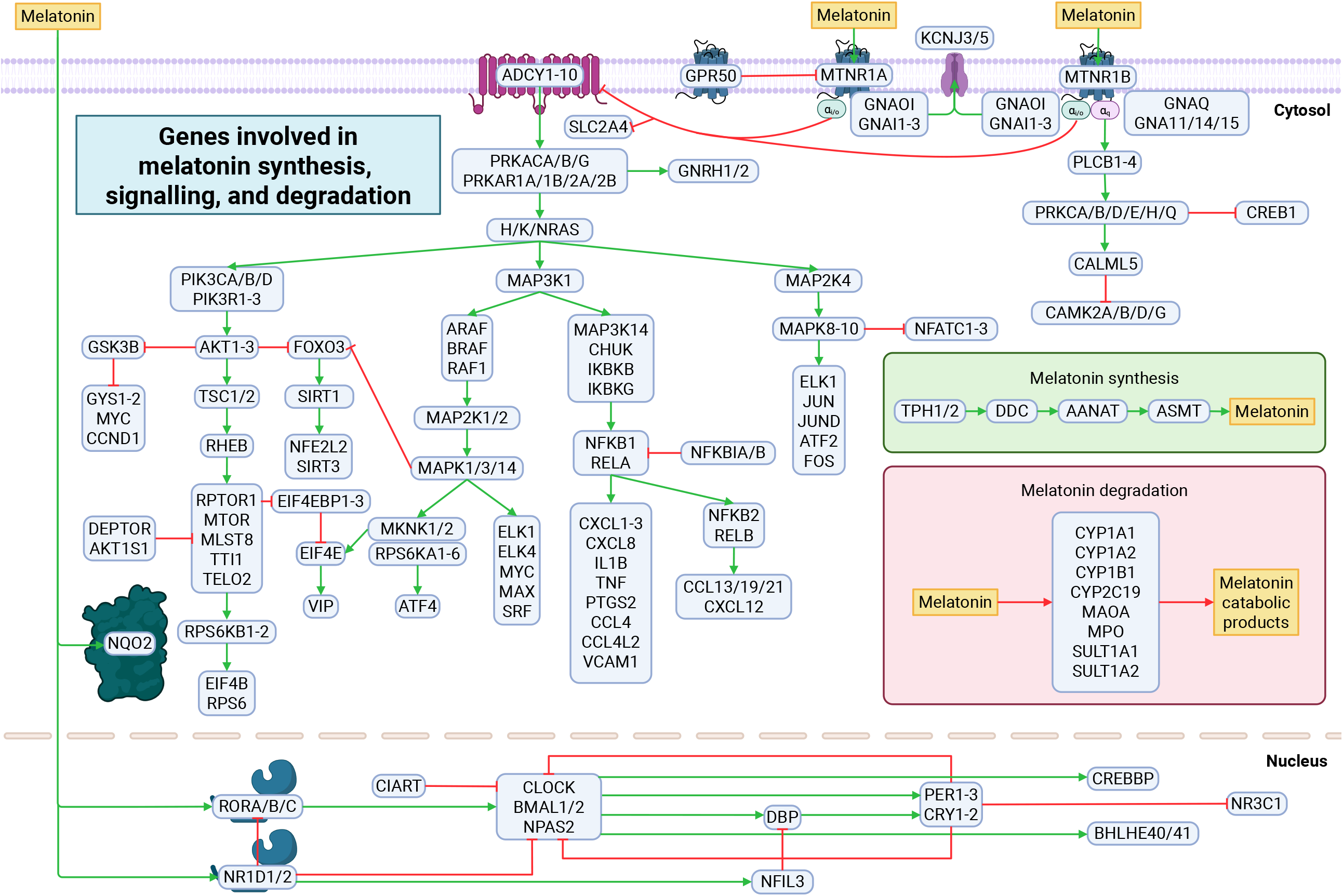
Genes involved in melatonin synthesis, signalling, and degradation. Melatonin is synthesised from serotonin via the enzymes AANAT and ASMT, primarily within the pineal gland. It then acts upon the cell-surface receptors MTNR1A and MTNR1B, nuclear receptors RORA/B/C and NR1D1/2, or the binding site NQO2. Binding to cell-surface receptors leads to activation of the PLCB1-4 gene pathway; and inhibition of ADCY1-10, which leads to the inhibition of the RAS/RAF/MAPK/ERK and PI3K/AKT/MTOR gene pathways, and activation of the SIRT1/FOXO3 pathway and GSK3B. Binding to nuclear receptors results in the rhythmic gene expression of core circadian clock proteins and transcription factors, which interact with one another via positive and negative feedback loops. Binding to NQO2 is thought to contribute to antioxidation and detoxification processes. [Created by authors with biorender.com]

A total of 181 genes found to be related to melatonin were identified (Table 1). These were then separated into “synthesis” (*TPH1/2, DDC, AANAT*, and *ASMT*), “degradation” (*CYP1A1*/*A2*/*B1*/*CA19, MAOA, MPO*, and *SULT1A1/2*), and “signalling” (i.e., melatonin-related genes not involved in the synthesis or degradation of melatonin). “Signalling” genes were then separated into “inhibitors” (i.e., genes that hinder melatonin signalling, or those whose expression is inhibited by melatonin) and “activators” (i.e., genes that enhance melatonin signalling such as receptors, or those whose expression is upregulated by melatonin). For example, melatonin activates the *PLCB* pathway via *MTNR1B*, which ultimately inhibits *CAMK2*; therefore, *MTNR1B* and *PLCB1-4* are labelled as “activators,” while *CAMK2A*/*B*/*D*/*G* are labelled as “inhibitors.” The full melatonin gene list can be found in Supplementary Table 2.

**Table 1:** Summary of melatonin gene list.

|  |  | Number of genes |
| --- | --- | --- |
| Signalling | Inhibitors | 110 |
|  | Activators | 58 |
| Synthesis |  | 5 |
| Degradation |  | 8 |
| **Total** |  | **181** |

### 3.2 Differential regulation of melatonin genes in people with bipolar disorder

Genes involved in the inhibition of melatonin signalling were found to be nominally altered in the BD post-mortem gene expression dataset with all sexes included (p = 0.0399, q = 0.1403; Table 2). The normalised enrichment score (NES) was positive, indicating overall upregulation of these genes (Table 2).

**Table 2:** Melatonin pathway gene expression from gene regulatory networks in people with bipolar disorder when compared to healthy controls.

| Sex | Melatonin gene pathways | ES | NES | p-value | q-value |
| --- | --- | --- | --- | --- | --- |
| All | Melatonin all genes | 0.259913 | 1.218872 | 0.092851 | 0.146606 |
|  | Melatonin signalling | 0.269811 | 1.258931 | 0.066621 | 0.140254 |
|  | Melatonin signalling activators | 0.212457 | 0.835406 | 0.776063 | 0.700208 |
|  | Melatonin signalling inhibitors | 0.312416 | 1.355442 | **0.039856** | 0.140254 |
|  | Melatonin degradation | -0.552180 | -1.099678 | 0.373603 | 0.471920 |
| Female | Melatonin all genes | -0.232726 | -1.068691 | 0.303193 | 0.353726 |
|  | Melatonin signalling | -0.234691 | -1.073567 | 0.291396 | 0.353726 |
|  | Melatonin signalling activators | 0.292640 | 1.132559 | 0.245732 | 0.353726 |
|  | Melatonin signalling inhibitors | -0.288312 | -1.228413 | 0.111447 | 0.353726 |
|  | Melatonin degradation | -0.504334 | -0.935858 | 0.565978 | 0.565978 |
| Male | Melatonin all genes | -0.282739 | -1.087871 | 0.267508 | 0.468139 |
|  | Melatonin signalling | -0.290515 | -1.113280 | 0.226789 | 0.468139 |
|  | Melatonin signalling activators | -0.345649 | -1.130808 | 0.249487 | 0.468139 |
|  | Melatonin signalling inhibitors | -0.264492 | -0.950058 | 0.564035 | 0.658040 |
|  | Melatonin degradation | 0.642688 | 1.135611 | 0.352053 | 0.492874 |

Abbreviations: ES = enrichment score; NES = normalised enrichment score.

TF-based GSEA revealed nominally significant downregulation of TFs related to the activation of melatonin signalling (p = 0.0483, q = 0.0552; Table 3) in the female-only BD post-mortem dataset. Significant downregulation of TFs related to melatonin signalling (p = 0.0055, q = 0.0043) and inhibition of melatonin signalling (p = 0.0055, q = 0.0043) were also found in the male-only BD post-mortem dataset (Table 3). The top ten genes and TFs modulated in people with BD when compared to HCs can be seen in Figure 3. Network-level targeting scores for melatonin pathway genes and transcription factors in people with BD compared with healthy controls are provided in Supplementary Tables 3 and 4.

**Table 3:** Melatonin pathway transcription factor expression from gene regulatory networks in people with bipolar disorder when compared to healthy controls.

| Sex | Melatonin TF pathways | ES | NES | p-value | q-value |
| --- | --- | --- | --- | --- | --- |
| All | Melatonin signalling | -0.376479 | -1.349456 | 0.105007 | 0.309686 |
|  | Melatonin signalling activators | -0.434801 | -1.237777 | 0.206457 | 0.309686 |
|  | Melatonin signalling inhibitors | -0.365951 | -1.113534 | 0.325463 | 0.325463 |
| Female | Melatonin signalling | -0.413200 | -1.464040 | 0.052407 | 0.055166 |
|  | Melatonin signalling activators | -0.538025 | -1.527795 | **0.048348** | 0.055166 |
|  | Melatonin signalling inhibitors | -0.333972 | -1.012314 | 0.439296 | 0.308278 |
| Male | Melatonin signalling | -0.477416 | -1.761388 | **0.005503** | **0.004274** |
|  | Melatonin signalling activators | -0.369348 | -1.066438 | 0.377778 | 0.132554 |
|  | Melatonin signalling inhibitors | -0.560702 | -1.732876 | **0.008120** | **0.004274** |

Abbreviations: ES = enrichment score; NES = normalised enrichment score; TF = transcription factor.

**Figure 3:**
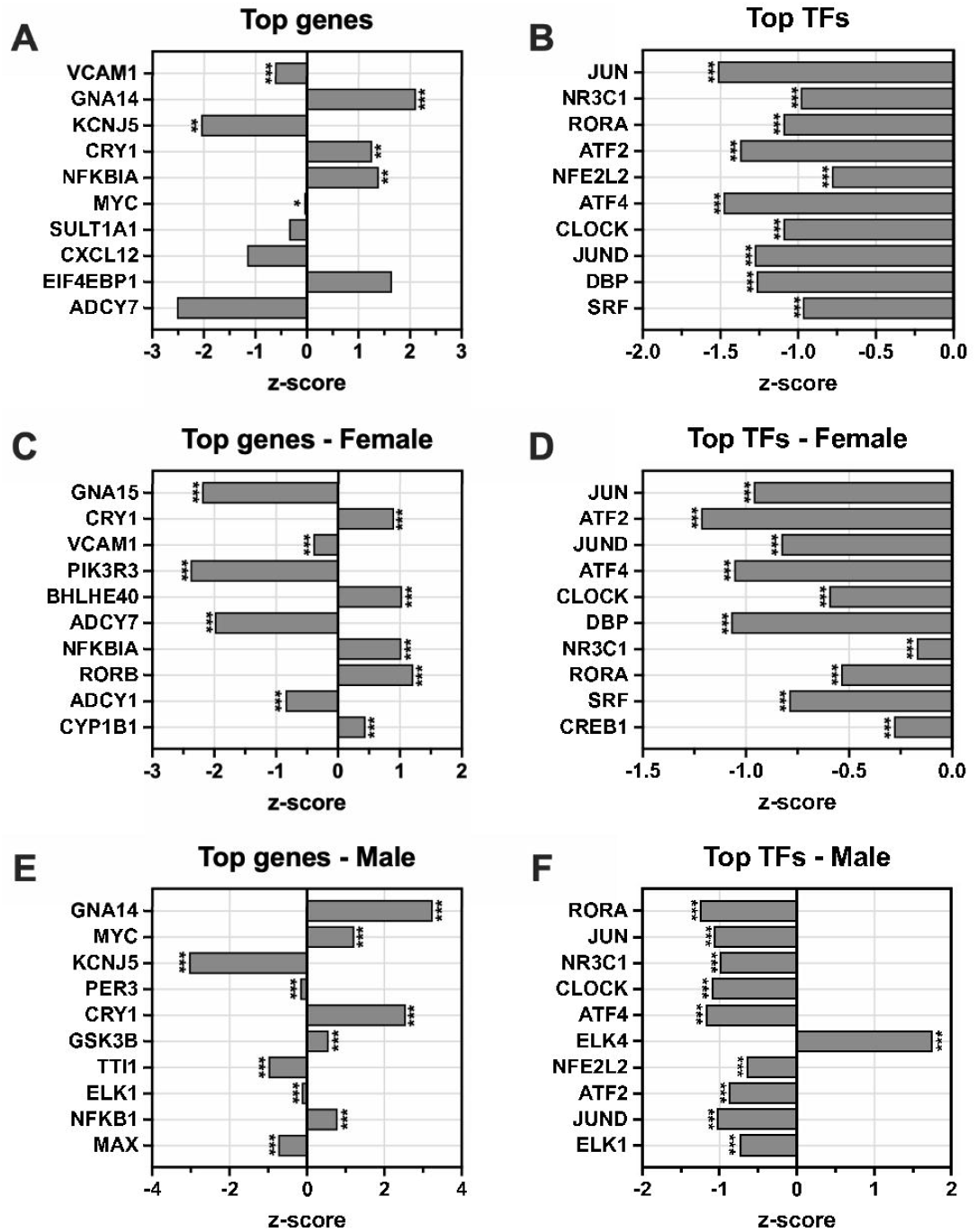
Top melatonin-related genes and transcription factors (TFs) differentially regulated between bipolar disorder patients and healthy controls. Top ten (ranked according to significance) genes ***(A)***, TFs ***(B)***, genes in females only ***(C)***, TFs in females only ***(D)***, genes in males only ***(E)***, and TFs in males only ***(F)*** from the melatonin gene list. Bars represent z-score; * q < 0.05, ** q < 0.01, *** q < 0.001.

### 3.3 Differential expression of melatonin genes by drugs used for bipolar disorder

The acute effects of lithium, quetiapine, valproate, and lamotrigine on melatonin-related genes were analysed by GSEA. These analyses unveiled only one significant result: a significant effect of valproate on genes involved in degradation of melatonin (p = 0.0003, q = 0.0026; Table 4). Valproate was shown to significantly upregulate these genes, as indicated by positive NES.

**Table 4:** Effects of commonly used medications for bipolar disorder on melatonin pathway gene expression.

|  |  | ES | NES | p-value | q-value |
| --- | --- | --- | --- | --- | --- |
| Lithium | Melatonin all genes | -0.301307 | -1.096476 | 0.279489 | 0.558977 |
|  | Melatonin signalling | -0.281008 | -1.018174 | 0.416447 | 0.666316 |
|  | Melatonin signalling activators | -0.388090 | -1.199307 | 0.202033 | 0.558977 |
|  | Melatonin signalling inhibitors | -0.206883 | -0.703698 | 0.952549 | 0.999772 |
|  | Melatonin degradation | -0.814531 | -1.416383 | 0.079383 | 0.558977 |
| **Quetiapine** | Melatonin all genes | 0.233966 | 0.937953 | 0.612688 | 0.994735 |
|  | Melatonin signalling | 0.208020 | 0.831219 | 0.857143 | 0.994735 |
|  | Melatonin signalling activators | 0.272096 | 0.917405 | 0.602989 | 0.994735 |
|  | Melatonin signalling inhibitors | 0.195728 | 0.734194 | 0.952812 | 0.994735 |
|  | Melatonin degradation | 0.776761 | 1.296761 | 0.163790 | 0.994735 |
| **Valproate** | Melatonin all genes | 0.314729 | 1.054478 | 0.380062 | 0.766172 |
|  | Melatonin signalling | 0.259843 | 0.868522 | 0.741636 | 0.766172 |
|  | Melatonin signalling activators | 0.256298 | 0.759618 | 0.831843 | 0.766172 |
|  | Melatonin signalling inhibitors | 0.266603 | 0.855901 | 0.732361 | 0.766172 |
|  | Melatonin degradation | 0.959691 | 1.661230 | **0.000347** | **0.002558** |
| **Lamotrigine** | Melatonin all genes | -0.156432 | -0.638202 | 0.999169 | 1 |
|  | Melatonin signalling | -0.159431 | -0.647747 | 0.998898 | 1 |
|  | Melatonin signalling activators | -0.222036 | -0.758627 | 0.880633 | 1 |
|  | Melatonin signalling inhibitors | 0.145015 | 0.520056 | 0.999675 | 1 |
|  | Melatonin degradation | 0.444832 | 0.731114 | 0.786813 | 1 |

Abbreviations: ES = enrichment score; NES = normalised enrichment score.

A chi-square test of independence showed a significant difference in the number of significantly altered melatonin-related genes between the four drug groups (χ^2^ (3, 597) = 124.0, p < 0.0001; Figure 4A). Quetiapine treatment caused the greatest number of significant alterations in melatonin-related gene expression, followed by valproate, lithium, and then lamotrigine (Figure 4A). A significant difference between groups was also found when the genes were further separated according to whether melatonin signalling was enhanced or opposed (χ^2^ (9, 597) = 126.3, p < 0.0001; Figure 4B). Overall, quetiapine treatment altered the expression of most of the investigated genes in favour of enhancing melatonin signalling, although the number of significantly-altered genes that enhanced melatonin signalling was only slightly greater than the number of significantly-altered genes that opposed melatonin signalling (Figure 4B). In valproate-treated cells, the number of significantly-altered genes that enhanced melatonin signalling and the number of significantly-altered genes that opposed melatonin signalling were equal (Figure 4B). Lithium significantly modified the expression of a small number of melatonin-related genes, and lamotrigine was not found to significantly alter the expression of any melatonin-related genes (Figure 4B). The top ten genes and TFs modulated in people with BD when compared to HCs is shown in Figure 4C-F. The effects of these mood stabilisers on the differential expression of all melatonin-related genes are presented in Supplementary Table 5.

**Figure 4:**
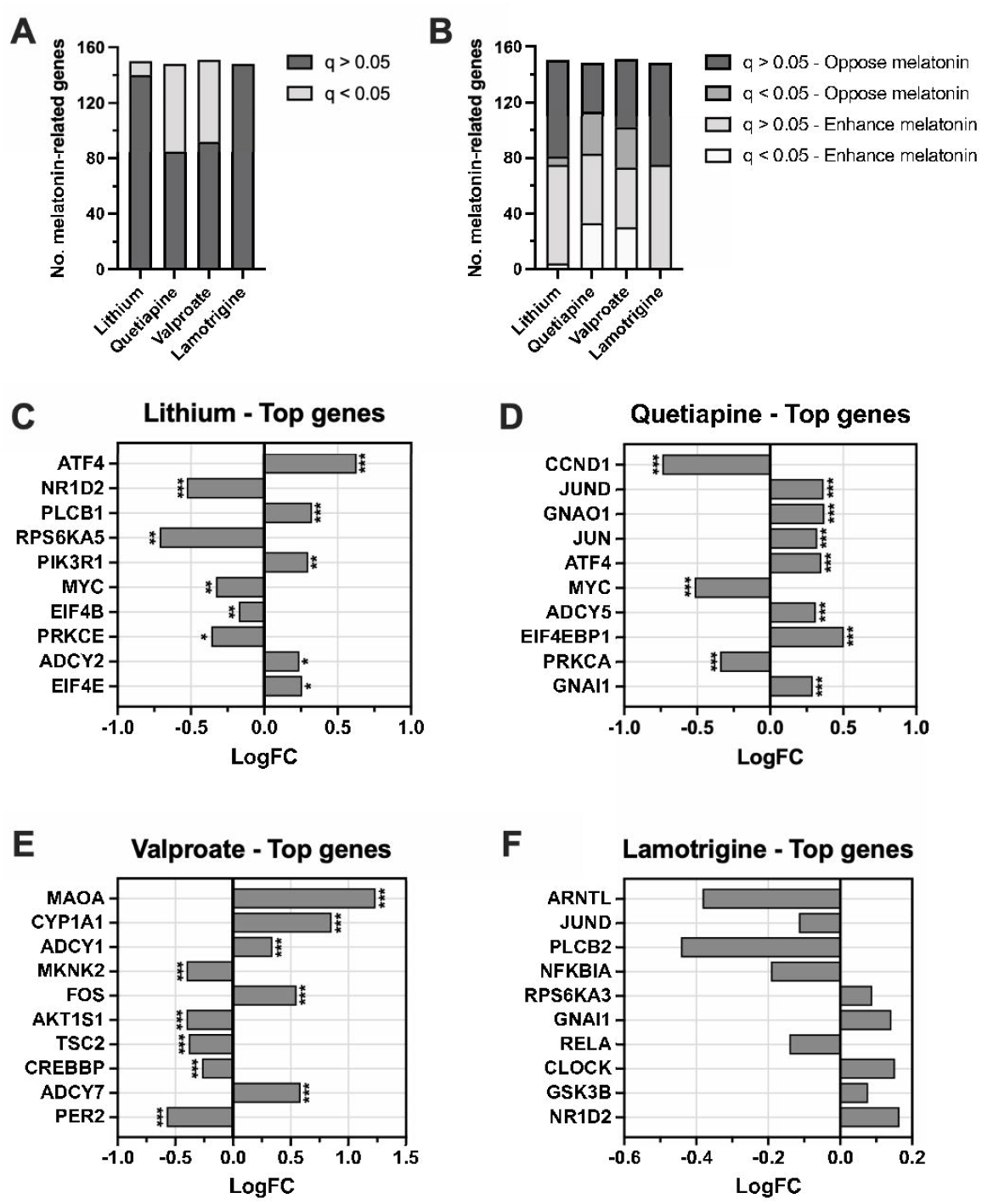
Effects of commonly used medications for bipolar disorder on differential expression of melatonin-related genes. Number of melatonin-related genes differentially expressed by lithium, quetiapine, valproate (VPA), and lamotrigine, separated by significance ***(A)*** and by significance and whether the genes oppose or enhance melatonin signalling ***(B)***. Top ten genes (ranked according to significance) from the melatonin gene list differentially expressed by lithium ***(C)***, quetiapine ***(D)***, VPA ***(E)***, and lamotrigine ***(F)***. Bars represent log fold change (LogFC); * q < 0.05, ** q < 0.01, *** q < 0.001.

### 3.4 Evaluating the effectiveness of repurposing melatonin modulators for bipolar disorder

Of the four melatonin agonists investigated, only the GR-135531 GRN was significantly dissimilar (as indicated by negative cosine; p < 0.0001, q < 0.0001) to BD-associated gene regulatory patterns (GRPs), suggesting it has potential as a drug repurposing candidate for BD (Table 5). The melatonin (p < 0.0001, q < 0.0001), ramelteon (p = 0.0006, q = 0.0016), and agomelatine (p < 0.0001, q < 0.0001) GRNs were significantly similar to BD-associated GRPs (Table 5).

**Table 5:**
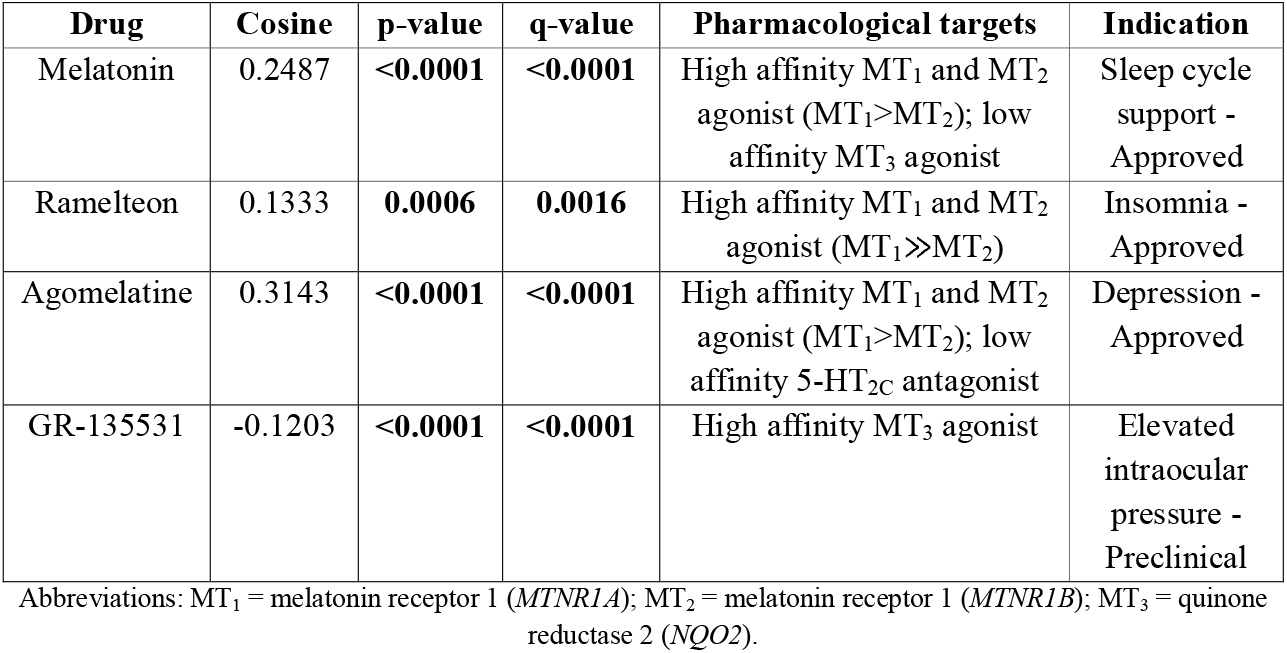
Effects of melatonin agonists on bipolar disorder-associated regulatory patterns.

When the effects of melatonin agonists on BD-associated GRPs were separated by sex, differences emerged. In females, the GR-135531 GRN was significantly dissimilar (p = 0.0038, q = 0.0154) to BD-associated GRPs, and the melatonin (p < 0.0001, q < 0.0001) and agomelatine (p < 0.0001, q < 0.0001) GRNs were found to be significantly similar (Table 6). In males, agomelatine (p = 0.0007, q = 0.0047) was significantly dissimilar, and ramelteon (p = 0.0242, q = 0.1047) was nominally significantly dissimilar to BD-associated GRPs (Table 6). GR-135531 GRN data were not available for males (Table 6). Full GRN results for all compounds in the full cohort, male-only, and female-only groups are listed in Supplementary Table 6.

**Table 6:** Effects of melatonin agonists on bipolar disorder-associated regulatory patterns – separated by sex.

|  | Female |  |  | Male |  |  |
| --- | --- | --- | --- | --- | --- | --- |
| Drug | Cosine | p-value | q-value | Cosine | p-value | q-value |
| Melatonin | 0.4352 | <0.0001 | <0.0001 | -0.0309 | 0.1452 | 0.4577 |
| Ramelteon | -0.0256 | 0.2993 | 0.7484 | -0.0448 | **0.0242** | 0.1047 |
| Agomelatine | 0.4957 | <0.0001 | <0.0001 | -0.1144 | **0.0007** | **0.0047** |
| GR-135531 | -0.0940 | **0.0038** | **0.0154** | N/A | N/A | N/A |

## 4 Discussion

The current study investigated the mechanistic role and therapeutic potential of melatonin in BD using transcriptomic analysis. To examine differences in the expression of genes implicated in melatonin synthesis, degradation, and signalling between people with BD and controls, we performed GSEA by comparing GRNs generated using RNAseq data derived from BD post-mortem brain samples against curated melatonin-related gene lists. Significant alterations in melatonin-related genes and TFs were observed in people with BD relative to controls, with notable sex-dependent differences. To explore whether currently approved mood stabilisers influence melatoninergic pathways, RNAseq data from NT2-N cells treated with lithium, lamotrigine, valproate, or quetiapine were analysed using the same GSEA approach to investigate whether current mood stabilisers for BD altered the expression of melatonin-related genes. We found that valproate significantly upregulated genes related to the degradation of melatonin. Finally, we compared BD GRPs to GRNs induced by melatonin receptor agonists (melatonin, agomelatine, ramelteon, and GR-135531) to evaluate the repurposing potential of these agents for BD. GR-135531 was identified as a promising repurposing candidate for people with BD, and agomelatine and ramelteon showed potential utility for males with BD only.

### 4.1 People with bipolar disorder demonstrate dysregulation of melatonin-related genes

An integral part of this study was the creation of a comprehensive list of genes related to melatonin signalling, synthesis, and degradation, and the subsequent classification of these genes as either “inhibitors” or “activators.” A small number of melatonin-related gene networks have been published; however, none adequately capture the widespread effects of melatonin on circadian entrainment, calcium homeostasis, metabolism, oxidative stress, and immune signalling. Typically, published gene networks are restricted to specific systems (e.g., apoptosis) or pathways (e.g., JAK/STAT signalling). In contrast, we took a novel approach and investigated all molecular and genetic pathways modulated by a single molecule, and used this to generate a comprehensive list that can be applied broadly by other researchers studying melatonin-related mechanisms.

This study utilised two types of GSEA: gene-based GSEA, which is used to examine downstream gene effects or pathway regulation; and TF-based GSEA, which is used to identify TFs that act as upstream drivers or key regulators. The gene-based GSEA results demonstrated a nominally significant upregulation of genes related to the inhibition of melatonin signalling in people with BD. Melatonin is a potent antioxidant, anti-inflammatory, and metabolic regulator^31,32^, and the signalling genes labelled as “inhibitors” were typically related to these functions. For instance, melatonin significantly downregulates mTOR complex 1 activity, leading to a reduction in glycolysis.^33^ The observed increase in genes related to inhibition of melatonin signalling may therefore reflect a state of metabolic dysregulation, inflammation, and/or oxidative stress in the BD dataset. People with BD frequently demonstrate increased markers of oxidative stress, chronic inflammation, and metabolic dysfunction when compared to healthy controls.^34–36^ Overall, these results suggest hypomelatoninaemia may directly contribute to biological pathophysiology of BD via an inability to suppress overactive gene pathways related to oxidative stress, inflammation, and metabolism. Therapeutic modulation of melatonin systems may therefore be effective in BD by targeting and normalising these dysregulated pathways.

While the TF-based GSEA did not reveal significant effects when both sexes were included, sex-stratified analyses uncovered interesting results. A prior analysis of the same post-mortem BD gene set with TF-based GSEA identified significant regulatory effects of TFs on immune response, energy metabolism, cell signalling, and cell adhesion.^13^ In this study, a nominally significant downregulation of TFs related to activation of melatonin signalling was noted in females with BD. This result appeared to be largely driven by the circadian genes *RORA* (which encodes the nuclear receptor RAR-related orphan receptor alpha (RORα)), *DBP, CLOCK*, and *BHLHE40*, all of which were found to be significantly downregulated. Activation of RORα by melatonin leads to a cascade of events contributing to circadian entrainment, including the activation and heterodimerisation of BMAL1 and CLOCK proteins.^37,38^ The BMAL1/CLOCK complex is a key modulator and stabiliser of the circadian clock, and regulates the expression of many other circadian genes, including *DBP, BHLHE40* and *BHLHE41*.^38^ Mutations in *CLOCK* and *RORA* have been implicated in the molecular pathophysiology of BD.^2,39,40^ Interestingly, males with BD also exhibited significant decreases in *CLOCK* and *RORA* gene expression, but the downregulation of TFs related to activation of melatonin signalling was not statistically significant. However, males with BD also exhibited significant downregulation of TFs related to the inhibition of melatonin signalling, likely driven by reductions in the expression of *JUN, JUND*, and *NR3C1*, all of which have well-established associations with BD.^13^

### 4.2 Valproate increased expression of genes related to the degradation of melatonin

Considering the complex relationship between melatonin and BD, examining the effects of BD medications on melatonin-related genes may be crucial for improving our understanding of the molecular mechanisms underpinning their clinical efficacy. Analysing psychotropic drug-induced gene expression changes may be key to elucidating their mechanisms of action, which is ultimately useful for predicting treatment response, identifying novel therapeutic targets, and tailoring pharmacological interventions.^41^

The results of this study demonstrated a significant effect of valproate on genes related to degradation of melatonin, including *MAOA* (which encodes monoamine oxidase A (MAO-A)), *CYP1A1*, and *SULT1A1*. Valproate has not been previously shown to upregulate MAO-A, which is the primary catabolic enzyme of not only melatonin, but also serotonin, dopamine, and noradrenaline. The augmenting effect of valproate on *SULT1A1* is also a novel finding. On the other hand, the effects of valproate on *CYP1A1* have been studied in detail in various cellular models, with all results showing consistent upregulation of the metabolic enzyme after valproate treatment.^42–44^ Increased degradation of melatonin may explain why valproate has previously been shown to blunt the nocturnal secretion of melatonin in healthy subjects.^45,46^ Unfortunately, no research has been conducted on the effects of valproate on melatonin secretion in psychiatric patients.

Although GSEA results were not significant for quetiapine, it is clear from individual gene expression analysis and Chi-squared tests that quetiapine has considerable effects on many melatonin-related genes. Of particular interest are quetiapine’s effects on circadian genes. Quetiapine was found to downregulate *BMAL1* and *BMAL2*, which, as previously mentioned, are essential for the stabilisation of the circadian clock; and upregulated the negative circadian regulatory genes *NR1D1* (which encodes nuclear receptor subfamily 1 group D member 1 (Rev-Erbα)), *CRY1, PER3*, and *NFIL3*. Activation of Rev-Erbα by melatonin leads to the recruitment of co-repressors that inhibit circadian gene expression.^37^ Furthermore, accumulations in CRY and PER proteins results in negative feedback that inhibits the transcriptional effects of CLOCK/BMAL1.^47^ Associations between mutations in *NR1D1, CRY*, and *PER* genes and BD have previously been identified.^47,48^ Overall, these results suggest that quetiapine may have a destabilising effect on circadian rhythm, which has implications for treatment.

Lithium and lamotrigine were found to have very little effect on melatonin-related genes. This supports clinical and preclinical work which have demonstrated mixed effects of lithium on melatonin secretion.^49–53^ The effects of lamotrigine on melatonin have not been published. Further investigation into the melatoninergic effects of mood stabilisers is therefore warranted.

### 4.3 GR-135531 may be a novel drug repurposing candidate for bipolar disorder

This study utilised a GRN analysis approach to measure the effectiveness of repurposing melatonin agonists for BD. This methodology was previously applied on the same post-mortem BD dataset to identify a number of novel drug repurposing candidates for BD, such as glutamine, iniparib, and alizapride.^13^ It was hypothesised that the melatonin GRN would exhibit significant dissimilarity to BD-associated GRPs, thereby indicating potential treatment effectiveness. Contrary to this hypothesis, the GRNs for melatonin, agomelatine, and ramelteon were found to be significantly similar to BD GRPs, suggesting these compounds may exacerbate, rather than alleviate, disease-related molecular changes in melatonin signalling in BD. In contrast, the dopamine GRN was found to be highly dissimilar from BD-associated GRPs, which may indicate the presence of reduced dopamine tone within the sample. Melatonin is known to exert potent antidopaminergic effects, as demonstrated in numerous preclinical studies.^54–62^ These results suggest that melatonin agonist usage may be contraindicated in people with BD who are experiencing depressive symptoms, who are frequently characterised by hypodopaminergic states. Additionally, GSEA revealed significant sex-dependent differences in response to melatonin agonists, suggesting that underlying sex-specific differences in dopamine or depressive symptomology may exist in this cohort.

GR-135531, a selective MT_3_ agonist, was also identified as a novel drug purposing candidate for BD. Preclinical evidence supports a role of GR-135531 in reducing intraocular pressure.^63^ MT_3_, which is a binding site on quinone reductase 2 (*NQO2*), is believed to mediate melatonin’s potent antioxidant activity.^64^ *NQO2* is expressed in multiple tissues, including the brain, heart, muscle, lung, intestine, and eye.^64^ Unlike the more extensively studied MT_1_ and MT_2_ receptors, MT_3_ represents an underexplored target for mood disorder research. As a result, no research on the preclinical effects of GR-135531 on behaviour or psychopathology has been published, which may indicate an avenue for future investigation.

### 4.4 Limitations

Despite several notable findings, this study has limitations that must be considered when interpreting results. First, clinical metadata for the BD post-mortem dataset were not available, including psychometric evaluations. As such, it is not known whether BD subjects were suicidal, or in euthymic, manic, mixed, or depressive mood states at the time of death, each of which may impact gene expression and treatment efficacy. In addition, these individuals were likely taking psychotropic medications, which could confound transcriptomic signatures. Further, only the DLPFC was sampled, yet gene expression profiles differ significantly across brain regions. Lastly, GR-135531 GRN data were not available for males, limiting sex-specific interpretation of this compound’s effects.

## 5 Conclusion

This study provides new evidence that dysregulation of melatonin-related genes may contribute to the pathophysiology of BD. People with BD may have a reduced capacity to suppress genes involved in oxidative stress, inflammation, and metabolic dysregulation, mechanisms that melatoninergic therapy may target. Furthermore, significant alterations were identified in TFs involved in the activation of melatonin signalling in females with BD, largely driven by downregulation of key clock genes. In contrast, males with BD showed significant downregulation of TFs associated with inhibition of melatonin signalling, primarily *JUN, JUND*, and *NR3C1*. GSEA and gene expression analysis demonstrated that valproate treatment significantly upregulated genes involved in melatonin degradation, potentially explaining its observed suppressive effects on melatonin levels in clinical studies. Quetiapine was also found to exert potentially destabilising effects on circadian genes. Finally, GR-135531 was identified as a potential drug repurposing candidate for BD. Results also suggest that melatonin treatment may be contraindicated in depressed people with BD due to its antidopaminergic properties. Collectively, the results of this study demonstrate that melatonin-related gene dysregulation may play a role in BD, underscore the need for sex-based analyses in psychiatric research, and position a number of melatoninergic agonists as potential therapeutics for BD.

## Acknowledgements

Heather K. Macpherson is supported by a Research Training Program stipend and tuition fee offset scholarship awarded by the University of Queensland. Michael Berk is supported by a NHMRC Leadership 3 Investigator grant (GNT2017131). Data were generated as part of the CommonMind Consortium supported by funding from Takeda Pharmaceuticals Company Limited, F. Hoffman-La Roche Ltd and NIH grants R01MH085542, R01MH093725, P50MH066392, P50MH080405, R01MH097276, RO1-MH-075916, P50M096891, P50MH084053S1, R37MH057881, AG02219, AG05138, MH06692, R01MH110921, R01MH109677, R01MH109897, U01MH103392, U01MH116442, project ZIC MH002903 and contract HHSN271201300031C through IRP NIMH.

